# MetaGenomic analysis of short and long reads

**DOI:** 10.1101/2020.03.13.991190

**Authors:** Felix Manske, Norbert Grundmann, Wojciech Makalowski

## Abstract

Identifying single organisms in environmental samples is one of the key tasks of metagenomics. During the last few years, third generation sequencing technologies have enabled researchers to sequence much longer molecules, but at the expense of sequencing accuracy. Thus, new algorithms needed to be developed to cope with this new type of data. With this in mind, we developed a tool called MetaG. An intuitive web interface makes the software accessible to a vast range of users, including those without extensive bioinformatic expertise. Evaluation of MetaG’s performance showed that it makes nearly perfect classifications of viral isolates using simulated short and long reads. MetaG also outperformed current state-of-the-art algorithms on data from targeted sequencing of the 16S and 28S rRNA genes. Since MetaG’s output is also supplemented with information about hosts and antibiotic resistances of pathogens, we expect it to be especially useful to the healthcare sector. Moreover, the outstanding accuracy of the taxonomic assignments will make MetaG a serious alternative for anyone working with metagenomic sequences. MetaG can be accessed at http://bioinformatics.uni-muenster.de/tools/metag/.

---

Microorganisms are ubiquitous: archeae and bacteria make up almost 14% of all biomass on this planet (humans: ca. 0.01%) as measured by mass of carbon (Bar-On et al. 2018). Consequentially, they are frequently used as markers in ecological studies to assess the status of habitats, e.g. (Bruni et al. 1997; Smith et al. 2015), or for bioremediation (Lovley 2003; Gihring et al. 2011). Additionally, the industry strives for discovering novel enzymes by metagenomics (Maurer 2004) and consequently 332 relevant enzymes were discovered between 2014 and 2017 (Berini et al. 2017). Microorganisms also impose a significant threat to human health (https://www.who.int/healthinfo/global_burden_disease/GHE_DthWBInc_Proj_2016-2060.xlsx?ua=1) and metagenomics is becoming increasingly popular among medical practitioners (Forbes et al. 2018). In the healthcare sector, metagenomics improves the diagnoses of patients with ambiguous symptoms (Wilson et al. 2014). Furthermore, it is critical for antibiotic resistance studies (Crofts et al. 2017).

To fuel discovery in healthcare, ecology and economy, downstream analyses need to be liberated. While sequencing technology is taking the first steps in this direction (see speed, price and portability of Nanopore devices (Loose 2017)), bioinformatics analyses are lagging behind: Metagenomic programs often suffer from excessive hardware requirements, e.g. using the native database of Kraken requires 70 GB of RAM (Wood and Salzberg 2014), and limited access. Additionally, the command line approach is a massive obstacle to less computer literate users. Established metagenomics programs often fail to handle the distinct sequencing error profiles of both short and long-read technologies (Santos et al. 2020). Although specific pipelines for Nanopore data already exist, they are mostly focused on specific usage cases (Santos et al. 2020).

MetaG was developed with these challenges in mind. An intuitive web interface allows researchers from different fields to run their analyses. The only requirement is an internet connection. However, for users working in more remote areas, we provide a local version of the software. Advanced users can customize their local runs to improve speed and/or accuracy to their needs. The source code of the core algorithm is publicly available at https://github.com/IOB-Muenster/MetaG/tree/master/metag_src to allow constant improvement by the community.

We compared MetaG to several state-of-the-art competitors using long and short reads of bacteria, archaea and fungi. In most cases, MetaG provided significantly improved classifications, especially at the species and strain levels. Additionally, short and long reads from three human pathogenic viral isolates were nearly perfectly classified. For each read type and database standard parameters are available.

## Results

### The core algorithm

MetaG uses LAST (Kiełbasa et al. 2011) to compare the user’s query reads to a database. Optionally, ambiguous alignments are filtered using LAST-SPLIT (Frith and Kawaguchi 2015). However, the analysis may also start from precomputed alignments given in MAF format (https://genome.ucsc.edu/FAQ/FAQformat.html#format5). At the time of writing, the employed databases are a customized version of RDP release 11.5 (Cole et al. 2014), a modified MTX database derived from the software METAXA 2.2 (Bengtsson et al. 2011) and the virus metadata resource (VMR) (https://ictv.global/vmr/) release MSL34 (version November 27) from the ICTV database (Walker et al. 2019) (see Supplemental Materials). These databases allow for analyses of bacterial, archaeal, fungal and viral samples. To assign the correct taxonomy to the reads, alignments are filtered based on user-given settings (see Figure 1). After removing alignments with too high e-values, the alignment scores per query read are analyzed (see Figure 1). Based on a relative threshold, only the alignments with an alignment score of at least T * maximum score, where T is a user-given fraction of one, are considered for downstream analysis.

**Figure 1:**
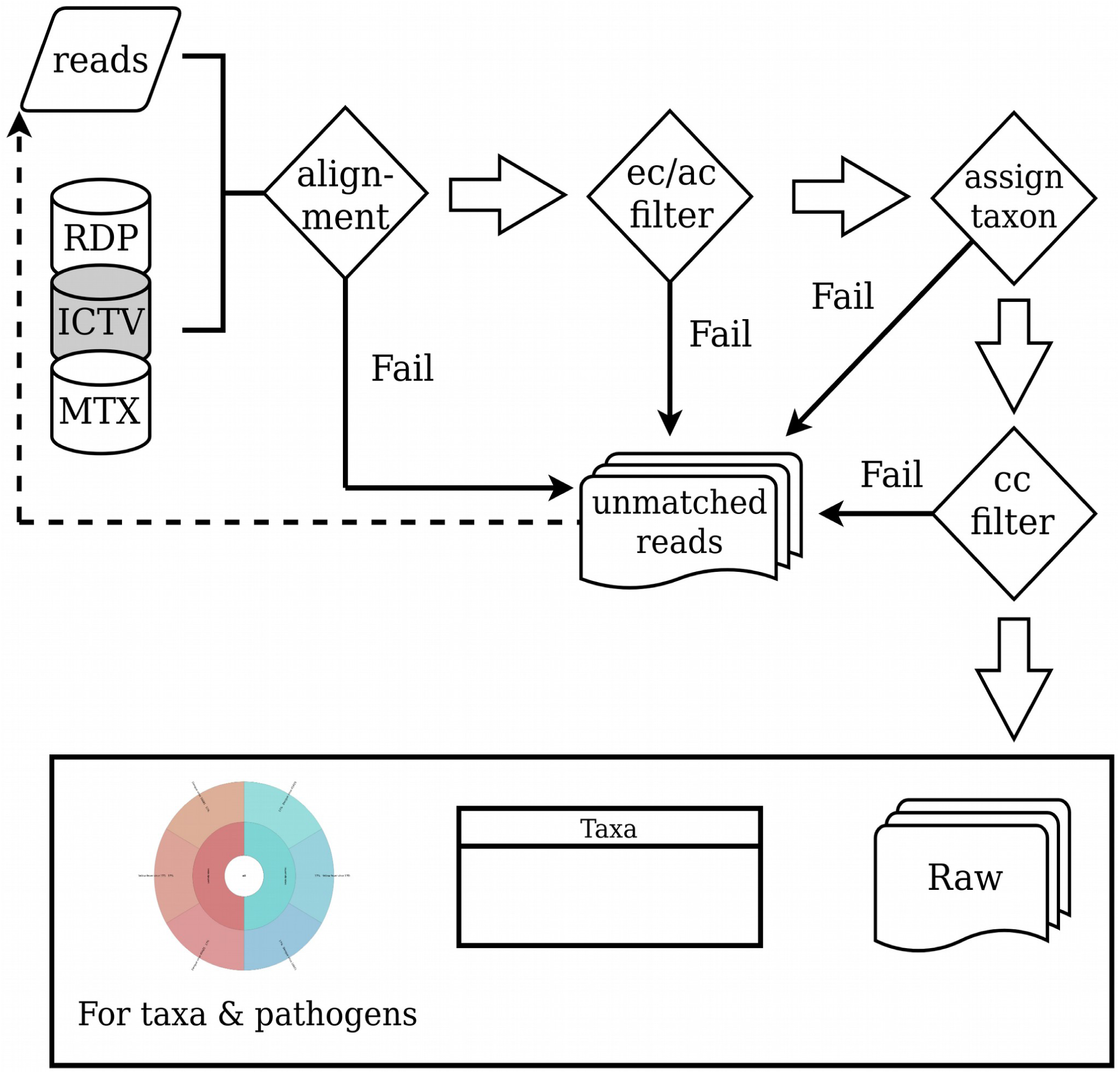
The core algorithm of MetaG exemplified by an analysis starting from fasta reads. Reads are first aligned to one (grey) of three available databases. The alignments are then filtered by an e-value cutoff (ec) and an alignment score cutoff (ac). Reads passing this filter are assigned with a taxonomic label and are subsequently filtered by a confidence cutoff (cc). Reads failing at any filter stage or failing the alignment itself are written to separate files for optional reanalysis. Taxon abundances of passing reads are presented as a table and as an interactive graph. Pathogen predictions and metadata are also shown as graphs. Raw data for plots and tables, the plots themselves and the taxonomic assignments on a per-read-basis are provided as raw data. This diagram was created with the online version of draw.io (https://www.draw.io/).

MetaG then assigns a taxonomic label to each read, starting at the broadest taxonomic level (see Figure 1). For this, the taxon given by most hits is used. The analysis progresses until multiple taxa are backed up by the same number of database hits of equal quality or a confidence threshold is violated (see Figure 1). The confidence threshold provides a measure of statistical support for the most abundant taxon that also reflects the quality of the underlying alignments (see Supplemental Materials). Reads failing to meet the assignment criteria are assigned to the unmatched class (see Figure 1).

MetaG provides information about taxonomic assignment at the level of individual reads and of the whole sample (see Figure 1). The numbers of input reads, matched reads and unassigned reads are supplemented with information about the run time and parameters. Users can access the number of reads assigned to an individual taxon and its average confidence value (see Figure 1). Unmatched read sequences at each taxonomic rank are written to separate files for further analysis. Reads that were filtered by the alignment process or e-value cutoff are written to a separate file (see Figure 1). This allows for reanalysis of these reads using a different database or different settings.

Found taxa and predicted pathogens are visualized in interactive graphs using KronaTools (Ondov et al. 2011) (see Figure 1). Where applicable, predicted pathogens are supplemented with host predictions and antibiotic resistance predictions. Bacterial and archaeal predictions are made at the species level, due to improved naming consistency across databases (data not shown). MetaG uses PATRIC (Wattam et al. 2017) to search for data on bacterial and archaeal pathogens. Viral pathogen data is derived from the same database as the taxonomy. Thus, predictions can be made at isolate level.

MetaG can be accessed via a web interface or installed locally. The latter version targets experienced users or those who want to perform analyses in remote areas without an internet connection. However, an installation script helps users to setup our program and most of its dependencies. The software was tested on Ubuntu 18.04.3 LTS, macOS 10.15.3 and FreeBSD 12.1. It depends on LAST 963, Perl v5.26.1 and KronaTools 2.7 or later versions. Users who are unfamiliar with a command line interface are encouraged to use the online version. Since calculations are performed on our servers, an internet connection is the only requirement. Due to its target audience, both versions of MetaG are supplemented with concise tutorials and standard parameters, which were specifically trained for short and long-read sequencing technologies. Reads can be supplied in fasta or fastq format.

### Benchmarking

After designing the algorithm we were curious to see how MetaG would perform on realistic sequencing data generated from representative approaches in metagenomics. For this, whole genome analyses of viruses and metaprofiling of 16S and 28S rRNA gene sequences were performed. Since the transition of short to long read sequencing data is one of the major challenges in the field of bioinformatics, both experiments used each of the two sequencing data types. Using metaprofiling data, we also wanted to assess MetaG’s performance in comparison to the current gold standard algorithms in the field.

### Whole genome analyses of viruses

We simulated MiSeqV3 sequencing of three pathogenic viral isolates at equal frequencies, including the generation of artifact reads. Using these data, MetaG made perfect classifications down to the isolate level. All reads which had been simulated as artifacts were not assigned (see Table 1). The found abundances of each virus were equal to the expected ones (see Table 1) and no additional isolates had been identified (data not shown).

**Table 1:** Percentage of reads assigned to an individual virus relative to the expectation in samples sequenced by the Illumina MiSeqV3 (*in silico*) and the MinION (*in vitro*). The UNMATCHED class represents reads which should not be assigned to any viral isolate.

| Taxon | MiSeqV3 | Nanopore |
| --- | --- | --- |
| Dengue virus 45AZ5 | 100 % | 101 % |
| Dengue virus 16681 | 100 % | 99 % |
| Yellow fever virus 17D | 100 % | 99 % |
| UNMATCHED | 100 % | 117 % |

Subsequently, we used genuine Nanopore data of the same isolates created by participants of the 2019 GRAID workshop in Sapporo, Japan (http://bioinformatics.uni-muenster.de/graid/education/workshops/hokkaido-2019-07/). Due to the nature of these data, the abundance of artifact reads was not available. We approximated the abundance of artifacts as the number of reads failing the initial LAST alignment. From the aligned reads, one-third was expected to come from each isolate (see Methods section). The observed abundance of the expected viruses deviated slightly from our expectations (see Table 1). This is partially due to the removal of reads during post-alignment processing in MetaG (see also Table 1). We also found 44 unexpected isolates, accounting for a total of 179 reads (data not shown).

### Simulated marker gene analyses

We were interested in how MetaG would perform in comparison to other established classifiers. Where possible, we wanted to analyze the differences between the RDP and MTX database for different algorithms (see Methods section). In a recent study of 16S rRNA data, Parallel-META v2.4.1 and QIIME v1.9.1 outperformed their competitors at the genus level (Escobar-Zepeda et al. 2018). For this reason, we chose to compare the performance of MetaG to these two programs. The RDP Classifier (Wang et al. 2007) is a lightweight program that can also be used online with pre-trained parameters. As we expected it to be optimized for the RDP database, we used it for our comparison. We employed the latest versions of the programs at the time of our analysis. QIIME 2 was run using its BLAST and Classifier workflow, respectively (see Methods section).

Matthew’s Correlation Coefficient (MCC) (Matthews 1975) (see Methods section) was used as an overall measure of the performance of MetaG and its competitors on MiSeq and Nanopore data. To use this measure, the exact origin of each read needed to be known (see Methods section). Thus, we simulated MiSeq and Nanopore sequencing of 27 reference 16S rRNA and 28S rRNA gene sequences from bacteria, archaea and fungi.

For the most part, the MCC was highest at the most general ranks and then decreased. After an increase, it reached a local maximum at the family or genus level, before dropping again (see Figures 2 and 3). From here on, this pattern will be called a wave-like pattern.

**Figure 2:**
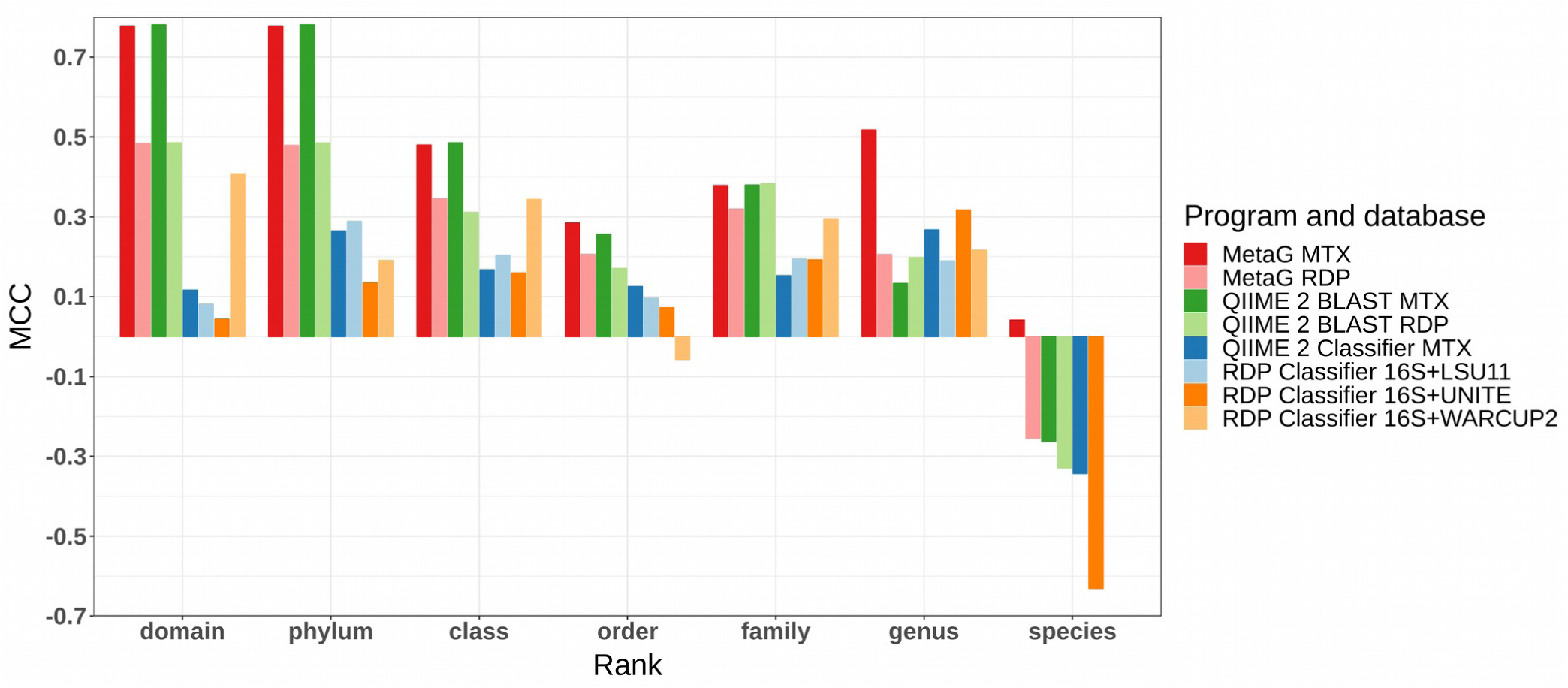
MCC from domain to species for classifiers using their chosen settings to analyze a sample containing bacteria, archaea and fungi. The sample was subject to simulated MiSeq sequencing. 16S is the 16S rRNA training set 16 of the RDP Classifier. The MCC for the RDP Classifier with the 16S database supplemented with LSU11 and WARCUP2, was undefined at species level. An MCC of one indicates perfect classifications. The MCC is minus one in case all taxonomic assignments were wrong.

**Figure 3:**
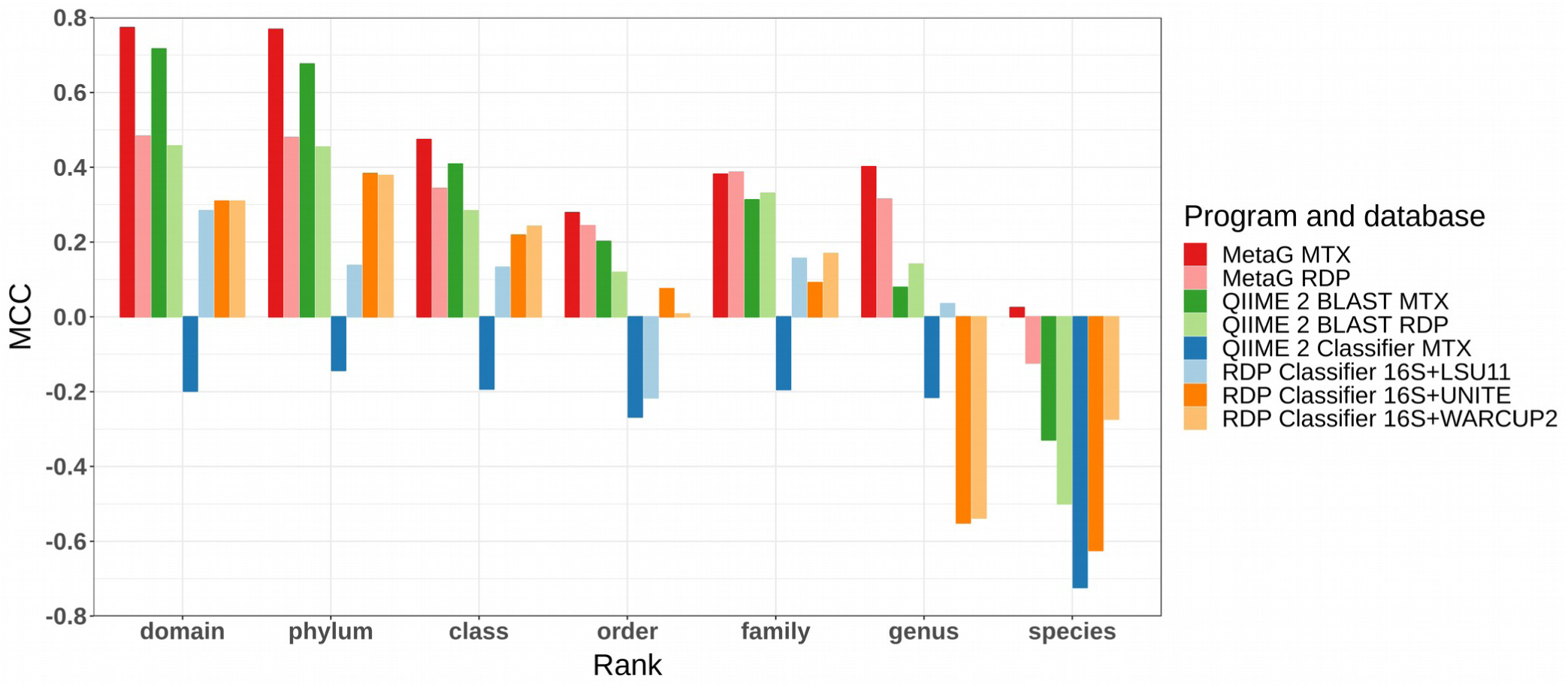
MCC from domain to species for classifiers using their chosen settings to analyze a sample containing bacteria, archaea and fungi. The sample was subject to simulated nanopore sequencing. 16S is the 16S rRNA training set 16 of the RDP Classifier. The MCC for the RDP Classifier with the 16S and LSU11 database was undefined at species level. An MCC of one indicates perfect classifications. The MCC is minus one in case all taxonomic assignments were wrong.

Using MiSeq data, the QIIME 2 Classifier and the RDP Classifier performed worst for most ranks up to the family level (see Figure 2). We could observe the same, but more pronounced, trend for Nanopore data. In this case, the performance was lowest up to the genus level (see Figure 3). Mostly, MetaG and QIIME 2 BLAST had the highest MCC up to the family level when using MiSeq reads. However, at the lower ranks, MetaG MTX was the most accurate program (see Figure 2). The results for our algorithm were even more promising for Nanopore data. This time, MetaG was always the best algorithm, as rated by MCC (see Figure 3). Using MTX, our program had the only positive MCC at species level, regardless of sequencing technology (see Figures 2 and 3). QIIME 2 BLAST MTX and the QIIME 2 Classifier performed substantially better using MiSeq data than Nanopore data (see Figures 2 and 3).

Naturally, the performance of algorithms was dependent on the database. This can be seen from the results of MetaG and QIIME 2 BLAST for which both RDP and MTX could be tested. When using MTX, algorithms generally performed better than when using RDP (see Figures 2 and 3). However, at family and genus level, this was inverse for QIIME 2 BLAST (see Figures 2 and 3). While these trends held true for both samples, we observed that RDP was slightly more beneficial than MTX for MetaG at the family level of Nanopore data.

### Analysis of a MinION mock sample

After the promising results from the previous analyses, we reanalyzed genuine Nanopore sequences from a bacterial community with a known abundance distribution. The data generated by Cuscó and coworkers had been previously analyzed by the authors using the Nanopore specific What’s in my Pot (WIMP) workflow (Cuscó et al. 2018; Juul et al. 2015).

We compared the expected bacterial abundances to the results by WIMP, MetaG and Parallel-META 3. The WIMP workflow underestimated the abundance of all expected taxa and overestimated the abundance of other taxa to a great extent (see Figure 4). Parallel-META 3 using its native database performed even worse, except in the case of *Staphylococcus aureus* (see Figure 4). Due to its performance on the simulated Nanopore data, MetaG was run using the MTX database. Notably, this workflow was always closest to the expected abundance of taxa with the exception of *Lactobacillus fermentum* and *Enterococcus faecalis* where the WIMP workflow was closest (see Figure 4).

**Figure 4:**
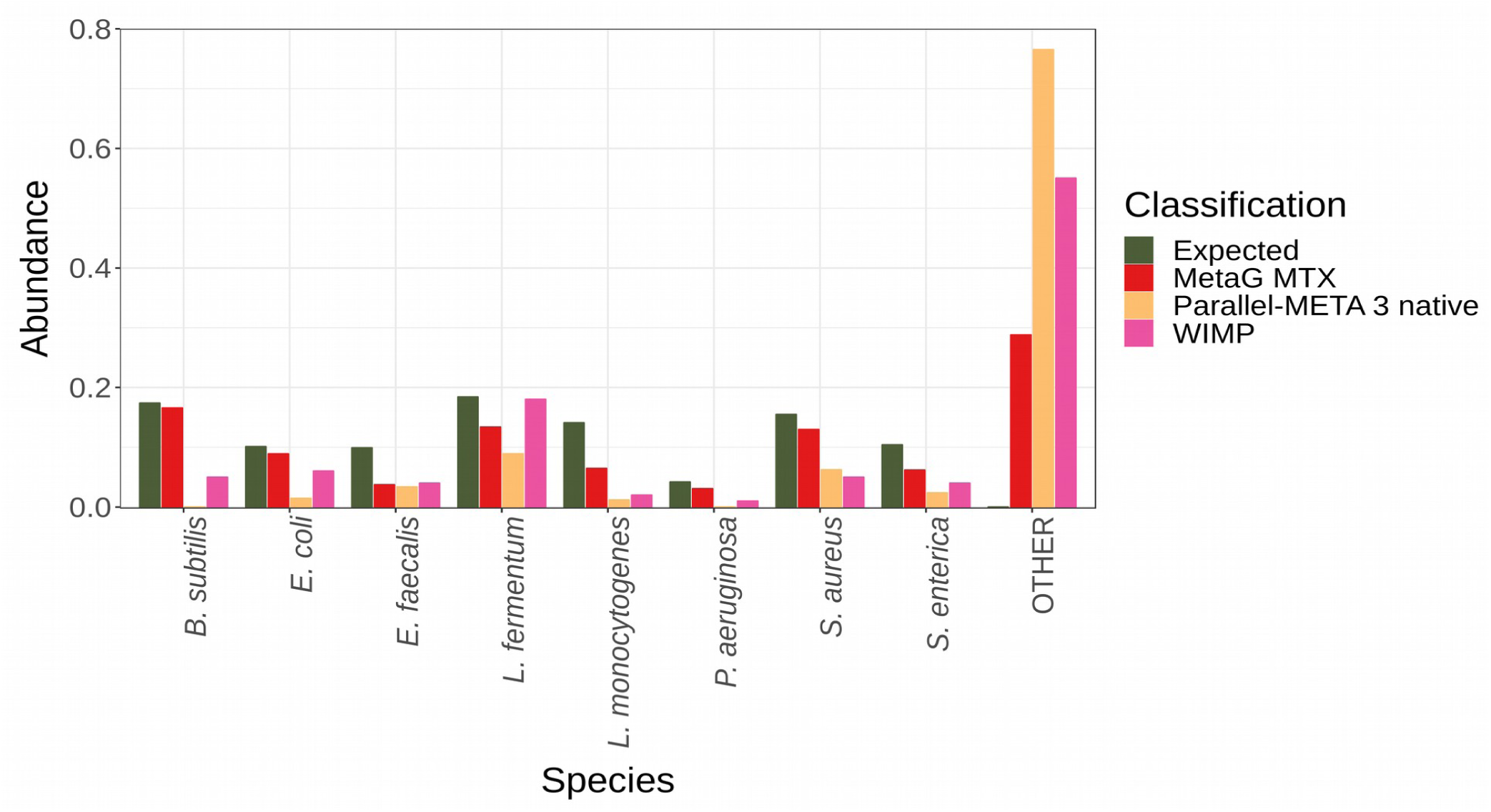
Bacterial classifications in the ZymoBIOMICS Microbial Community DNA Standard D6306 sequenced by the MinION. Relative abundances given by MetaG MTX, Parallel-META 3 and the WIMP workflow of Cuscó and coworkers (Cuscó et al. 2018) are compared to the expected classifications for the mock sample (https://files.zymoresearch.com/protocols/_d6305_d6306_zymobiomics_microbial_community_dna_standard.pdf).The OTHER class sums up all taxa which were not expected.

## Discussion

When starting the development of MetaG, we recognized that especially usability and portability of algorithms in the field were in need of improvement. Accordingly, we set the following goals for ourselves. Firstly, in order to fuel biological discoveries, we aimed to create a software which is open to as many researchers as possible. Secondly, by allowing free online analysis on our servers, we enable researchers with tight budgets or older hardware and bioinformatic novices to run their analyses. Researchers who are familiar with command line, use state-of-the-art hardware or who are working in more remote areas without an internet connection are encouraged to try the local version of MetaG. An installation script makes the setup as simple as the installation of any regular software. While the online version is virtually independent of the operating system, for the local version, we took great care to support most of the operating systems used in bioinformatics. The installer was tested on macOS 10.15.3, Ubuntu 18.04.3 LTS and FreeBSD 12.1. Running the local version also allows for speeding up the process by assigning more computational resources. Additionally, our program allows for a high level of customization. Unlike Parallel-META 3 (see Methods section), MetaG allows for analysis using any correctly formatted database. Parameters for custom databases or future sequencing technologies can be obtained by using our training routine (see Methods section). Users are encouraged to improve the algorithm or even run their own publicly available MetaG server.

We aimed to create a software that is both easy to use and shows high performance. By using simulations and real sequencing data, we have shown that MetaG is capable of performing very precise assignments. Whole genome sequencing of three human pathogenic viruses showed perfect classifications for MiSeq data and near-perfect classifications for MinION data (see Table 1). Comparing species identifications of *in vitro* 16S rRNA gene reads, we demonstrated that MetaG was overall ahead of the competition by predicting the most realistic species abundances (see Figure 4). Strikingly, the study, which produced the *in vitro* reads (Cuscó et al. 2018), used the WIMP workflow which was published specifically for the MinION (Juul et al. 2015). Thus, MetaG is powerful enough to beat algorithms at their own game.

When analyzing bacteria, archaea and fungi, the same trend was apparent and especially pronounced at the lower ranks: MetaG using MTX was the only software with a positive MCC at the species level (see Figures 2 and 3). Since the pathogenicity of *Escherichia coli* varies within the species (Johnson 2002), precise identifications at the strain level are desirable for most applications of metagenomics.

In most cases, the alignment-based algorithms, MetaG and QIIME2 BLAST, displayed a better performance than the kmer-based approaches, namely the QIIME2 and RDP classifiers (see Figures 2 and 3). This is in line with previously identified problems with the kmer-based approaches (Gao et al. 2017). Additionally, the used databases also influenced the outcomes. In most cases, using MTX was preferable over RDP (see Figures 2 and 3). This is likely due to the high level of manual curation of MTX (Bengtsson-Palme et al. 2015).

Sequencing technology was also a major factor impacting the algorithm performance. The QIIME2 Classifier and QIIME2 BLAST MTX, for example, performed better when using MiSeq data (see Figures 2 and 3). This indicates that some algorithms could not handle the trade-off between error rate and read length. MetaG’s results were consistently showing high performance (see Figures 2 and 3). This can be attributed to the training of parameters for each sequencing technology and database (see Methods section).

The observed wave-like MCC pattern was likely due to naming inconsistencies between the reference taxonomy from NCBI and the taxonomies in MetaG’s different reference databases. In an earlier study, the authors only focused on phylum and genus identifications to avoid this artifact (Lindgreen et al. 2016). The differences between the algorithms cannot be explained by these inconsistencies. The general drop in MCC towards the species level was real and related to missing resolutions of the marker genes (Knight et al. 2018). This can also be seen from our whole-genome analyses of viruses; they have shown that the vast majority of reads were assigned to the correct isolate (see Table 1). In contrast, the species identifications for metaprofiling of bacteria, archaea and fungi were less definitive (see Figures 2 and 3). However, due to speed and ease of use, metaprofiling is still popular (Knight et al. 2018).

In order to make high performance available to the community, software development must not stop at creating a precise algorithm. Rather, the user must also be able to use the software accordingly. We supplemented MetaG with a concise manual and strong standard settings for short and long reads and the three different databases. As our analyses have shown, these are usually sufficient to allow users to get to the full potential of MetaG, regardless of the sequencing approach. However, the standard settings also provide an excellent starting point to adjust the parameters to the individual experiment.

In summary, we have reached our goal of designing a portable, easy to use software with outstanding performance. In the light of our results, MetaG will improve analysis in areas such as medical metagenomics and ecologically and economically motivated metagenomics. By identifying or predicting pathogens and antibiotic resistances, MetaG provides essential features, especially for the healthcare sector. By design, our software supports adaptations to other current or future databases and sequencing technologies. Thus, we invite everyone to use MetaG for their individual projects and to help improve the software by giving feedback or modifying the source code.

## Methods

### Obtaining standard parameters

To achieve the best performance, the parameters of MetaG were trained. To accomplish this, a short- and a long-read sample were analyzed using a single database at a time. By using LAST-TRAIN (Hamada et al. 2017), we obtained the optimal alignment parameters for LAST. The optimal values for LAST-SPLIT filtering, e-value, alignment score and confidence cutoff were determined manually with the help of custom semi-automatic training scripts (see Supplemental Materials). The standard parameters are accessible from https://github.com/IOB-Muenster/MetaG/tree/master/metag_src/install/files as config files for MetaG.

### Whole genome analyses of viruses

In July 2019, participants of the GRAID workshop in Sapporo, Japan sequenced Yellow fever virus 17D and Dengue virus type 1 and 2 using the MinION flowcell FLO-MIN107 (R9.5). Each of the viruses was exclusive to four of twelve barcoded samples. We took 4,000 debarcoded pass reads from each sample. The portions were pooled and analyzed in MetaG. We inspected the numbers of reads matching to the expected taxa and to the unmatched class. Due to the nature of the sample, we had to approximate the expected number of unmatched reads (775 or about 2 percent of the reads) as the observed abundance of reads without any alignment.

To get realistic short reads, we simulated MiSeqV3 sequencing using ART version 2.5.8 (Huang et al. 2012). Three isolate sequences were obtained from GenBank (Benson et al. 2018): Yellow fever virus 17D (FJ654700.1), Dengue viruses type 1 (M87512.1) and type 2 (M29095.1). Three different samples were created for each of the viral isolates. Each sample contained 16,000 reads with a length of 250 bases. The following command was executed: art_illumina -nf 0 -ss MSv3 -amp -na -q -i [sample.fasta] -l 250 -f 16000 -o [out.fastq]. The individual fastq files were transformed to fasta and the first 960 reads of each sample were shuffled using a customized script (initial source: https://github.com/Ales-ibt/Metagenomic-benchmark/blob/master/bin/16SrRNAamplicon/shuffled_fasta.pl) to simulate sequencing artifacts. Next, the three samples were pooled. The number of reads in the pooled *in vitro* MinION and *in silico* MiSeq samples was 48,000, each. Analyses were performed using MetaG. The abundances for the three viral isolates and the unmatched, i.e. shuffled, reads were compared to what was expected. ICTV VMR release MSL34 (version November 27) lists Dengue virus 45AZ5 and 16681 as virus isolate designations of the commonly used virus names Dengue virus type 1 and 2, respectively. We acknowledged this by setting the expected virus names to the viral isolate designations given by ICTV.

### Simulated sequencing of marker genes

We obtained 27 reference 16S or 28S rRNA gene sequences from bacteria, archaea and fungi from the NCBI Nucleotide database (Sayers et al. 2019) (see Table 2). The sequences showed different degrees of relatedness. Three of the sequences belonged to fungi or archaea, respectively. One sequence stemmed from the bacterium *Xylanibacillus composti* K13, which had only recently been described as *genus novum* and *species novum* (Kukolya et al. 2018).

**Table 2:**
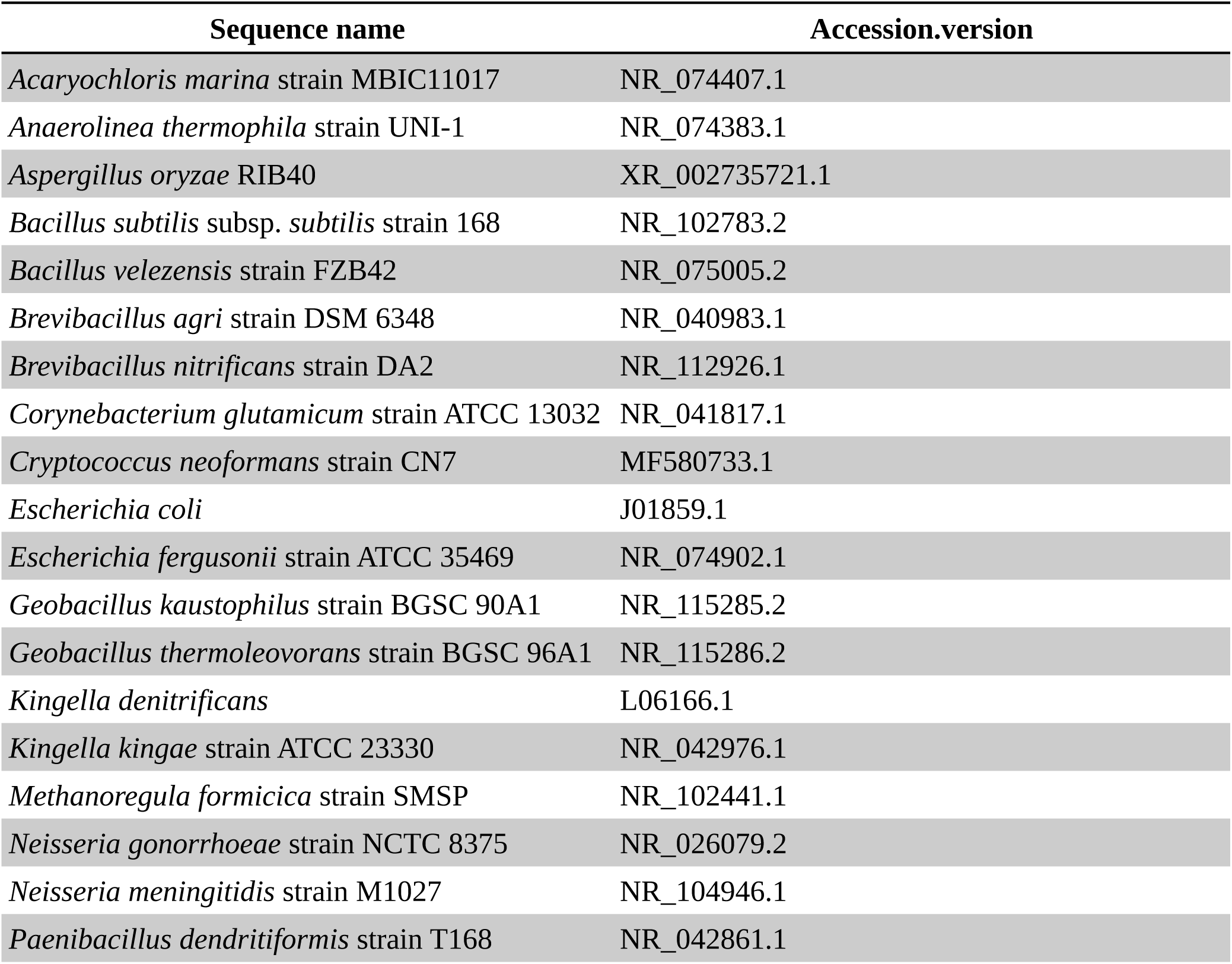
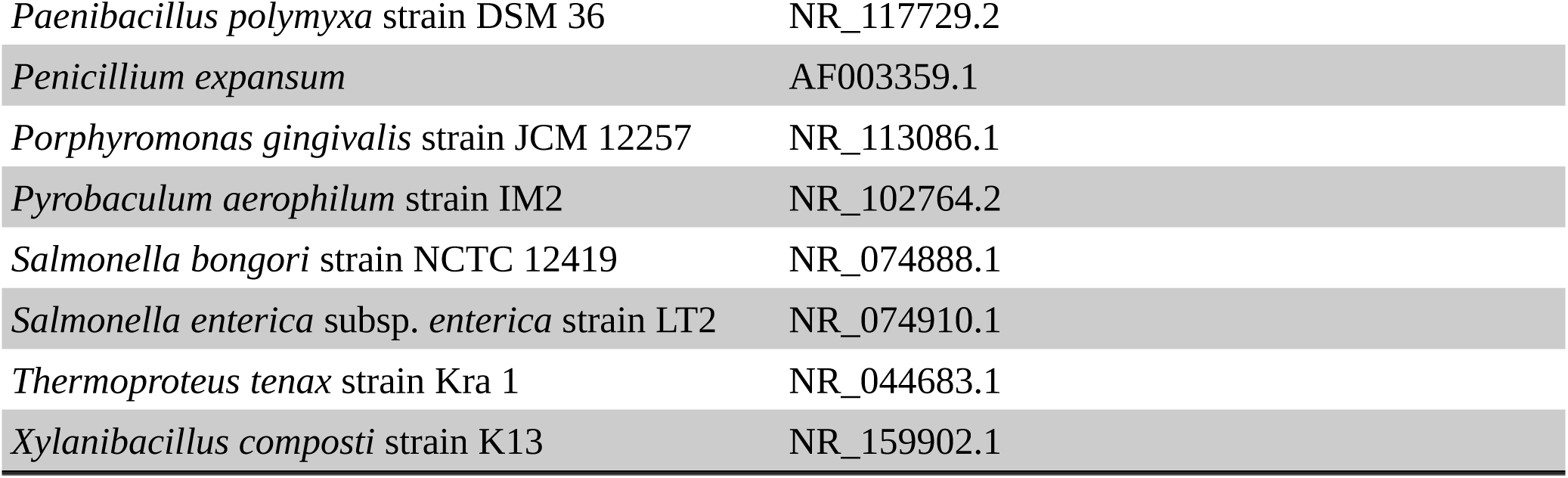
Accession and version of the sequences retrieved from the NCBI Nucleotide database.

| Sequence name | Accession.version |
| --- | --- |
| *Acaryochloris marina* strain MBIC11017 | NR_074407.1 |
| *Anaerolinea thermophila* strain UNI-1 | NR_074383.1 |
| *Aspergillus oryzae* RIB40 | XR_002735721.1 |
| *Bacillus subtilis* subsp. *subtilis* strain 168 | NR_102783.2 |
| *Bacillus velezensis* strain FZB42 | NR_075005.2 |
| *Brevibacillus agri* strain DSM 6348 | NR_040983.1 |
| *Brevibacillus nitrificans* strain DA2 | NR_112926.1 |
| *Corynebacterium glutamicum* strain ATCC 13032 | NR_041817.1 |
| *Cryptococcus neoformans* strain CN7 | MF580733.1 |
| *Escherichia coli* | J01859.1 |
| *Escherichia fergusonii* strain ATCC 35469 | NR_074902.1 |
| *Geobacillus kaustophilus* strain BGSC 90A1 | NR_115285.2 |
| *Geobacillus thermoleovorans* strain BGSC 96A1 | NR_115286.2 |
| *Kingella denitrificans* | L06166.1 |
| *Kingella kingae* strain ATCC 23330 | NR_042976.1 |
| *Methanoregula formicica* strain SMSP | NR_102441.1 |
| *Neisseria gonorrhoeae* strain NCTC 8375 | NR_026079.2 |
| *Neisseria meningitidis* strain M1027 | NR_104946.1 |
| *Paenibacillus dendritiformis* strain T168 | NR_042861.1 |
| *Paenibacillus polymyxa* strain DSM 36 | NR_117729.2 |
| *Penicillium expansum* | AF003359.1 |
| *Porphyromonas gingivalis* strain JCM 12257 | NR_113086.1 |
| *Pyrobaculum aerophilum* strain IM2 | NR_102764.2 |
| *Salmonella bongori* strain NCTC 12419 | NR_074888.1 |
| *Salmonella enterica* subsp. *enterica* strain LT2 | NR_074910.1 |
| *Thermoproteus tenax* strain Kra 1 | NR_044683.1 |
| *Xylanibacillus composti* strain K13 | NR_159902.1 |

Subsequently, we simulated 400 MiSeq reads per sequence as described for the viral sequences. The fastq files were transformed to fasta and the first 24 reads of each reference sequence were shuffled to simulate sequencing artifacts (see previous section). All reads were pooled and used for subsequent analysis.

We simulated the results of 2D MinION sequencing with a flowcell version of R9 by using NanoSim-H version 1.1.0.4 (Yang et al. 2017; Břinda and Yang). According to the shortest sequence obtained from NCBI, the maximum read length was set to 1,415 bases. Again, 400 reads for each sequence were produced by executing the following command: nanosim-h -p ‘ecoli_R9_2D’ -n 400 --circular --max-len 1415. Nanosim-H automatically created 24 unalignable reads for each sequence. This corresponds to the custom shuffling approach when using ART. The resulting fasta files were pooled to a single sample. The reads were then analyzed using several programs (see next section).

To evaluate the results, the classifications of the individual reads were compared to the expectation derived from the NCBI Taxonomy database (Sayers et al. 2019). During our preliminary analysis, we noted that strain assignments to metaprofiling data were somewhat arbitrary. This was in line with previous findings (Knight et al. 2018). Accordingly, the lower limit for the evaluation of all programs was the species level.

The classifications were evaluated as follows. If a non-shuffled read was assigned to the correct taxon, it was a true positive (TP), otherwise it was a false positive (FP). If it could not be assigned, it was a false negative (FN). A shuffled read that was assigned to a taxon was a false positive (FP), otherwise it was a true negative (TN). Using the above classes we calculated the Matthew’s Correlation Coefficient (MCC) for all analyses as an overall performance indicator (Matthews 1975).

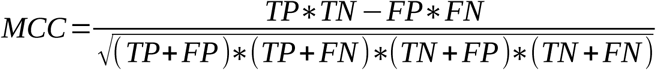

Plots were created using R 3.4.4 (R Core Team 2018) and the packages tidyverse 1.2.1 (Wickham 2017) and reshape2 1.4.3 (Wickham 2007).

### Performance evaluation of taxonomic assignments

We chose to compare MetaG to the topical versions of the RDP Classifier, Parallel-META and QIIME. The latter two programs had previously shown excellent performance at genus level (Escobar-Zepeda et al. 2018). At the time of the study, Parallel-META v2.4.1 could be run with several databases using a patch (Escobar-Zepeda et al. 2018). However, this was not possible using Parallel-META 3 (Jing et al. 2017), which was the newest version at the time of our analysis. Therefore, we could only use Parallel-META 3’s custom database. As our sample contained 28S rRNA sequences, which were not supported by Parallel-META 3’s database, we only used the program on the bacterial mock sample (see next section).

We used the command line version of QIIME 2 (Bolyen et al. 2018) version 2018.11.0. Samples and databases were imported using standard settings. Each sample was considered to be demultiplexed. No quality filtering was performed, since the data was in fasta and not in fastq format. We skipped the chimera filtering, as the simulations did not involve any artificial PCR. Following advice from QIIME2 support (https://forum.qiime2.org/t/analysis-of-fastq-files/3177), we did not perform any denoising for the nanopore data. To get comparable results, denoising was also skipped for the MiSeq data.

To obtain the correct input format (QIIME2 term: artifact) for the analyses, the samples had to be dereplicated. However, this resulted in a loss of some sequence IDs, which would have complicated our downstream benchmarking. Thus, we replaced the sequences inside the FeatureData[Sequence] artifact with the full set of sequences. We took care to adapt the sequences to the QIIME2 format, e.g. the maximum amount of characters per line was 80.

Subsequently, we analyzed the sequences using two different approaches. The first used a BLAST+ (Camacho et al. 2009) implementation of QIIME2 with standard settings on the RDP and MTX database:

> qiime feature-classifier classify-consensus-blast
>
> --i-query [FeatureData[Sequence]]
>
> --i-reference-reads [FeatureData[Sequence]]
>
> --i-reference-taxonomy[FeatureData[Taxonomy]]

The other approach used a QIIME2 implementation of a Naїve Bayesian Classifier ve Bayesian Classifier (Pedregosa et al. 2011; Bokulich et al. 2018). First, the classifier was trained on RDP and MTX. However, in the case of the former database, this was not successful; possibly due to the number of database records. After training, query reads were classified using standard settings and the MTX database:

> qiime feature-classifier classify-sklearn
>
> --i-reads [FeatureData[Sequence]]
>
> --i-classifier [training_profile]

Both QIIME2 implementations provided taxa with confidence levels. We examined the output for both implementations and the used databases with the confidence thresholds 0, 0.5 and 1. We chose to filter the taxa by the confidence cutoff that provided the highest overall performance for each database and sequencing technology, as indicated by the MCC (optimal threshold). If two confidence thresholds yielded similar performance, the stricter one was chosen. For the BLAST+ and the classifier implementations, the chosen confidence cutoff was 0.5, regardless of sequencing technology and database (data not shown). These thresholds were applied to both simulated Nanopore and MiSeq data.

At the time of analysis, the online implementation of the RDP Classifier contained four databases: *16S rRNA training set 16* (https://rdp.cme.msu.edu/classifier/classifier.jsp), *Fungal LSU training set 11 (Liu et al. 2012), Warcup Fungal ITS trainset 2 (Deshpande et al. 2016)* and the *UNITE Fungal ITS trainset 07-04-2014 (Wang and Cole 2014)*. The former database was a 16S rRNA gene database, the others focused on fungal sequences. Thus, fungal and bacterial/archaeal sequences had to be analyzed separately. The results obtained for MiSeq and Nanopore data using the *16S rRNA training set 16* were merged with the results obtained by using each of the fungal databases and the respective confidence cutoff.

The optimal confidence thresholds were determined as described for QIIME2. The optimal thresholds for the 16S rRNA database merged with *Fungal LSU training set 11, Warcup Fungal ITS trainset 2* and *UNITE Fungal ITS trainset 07-04-2014* for Nanopore and MiSeq sequencing (square brackets) were 1.0 [0.5], 0.5 [1.0] and 0.5 [0.5], respectively.

### Analysis of a MinION mock sample

Cuscó and coworkers sequenced the ZymoBIOMICS Microbial Community DNA Standard D6306 using nanopore chemistry R9.4.1 and 1D reads (Cuscó et al. 2018). We obtained the reads from the Sequence Read Archive (SRA) (Leinonen et al. 2011) using run ID SRR8029984 (Cuscó et al. 2018). Cuscó et al. classified the 16S rRNA gene sequences by using the What’s in my Pot (WIMP) (Juul et al. 2015) workflow in conjunction with the NCBI database. They compared the results to the manufacturer’s expectation (https://files.zymoresearch.com/protocols/_d6305_d6306_zymobiomics_microbial_community_dna_standard.pdf) (Cuscó et al. 2018). In line with the study, we applied 16S rRNA gene-based workflows: we chose to use MetaG MTX on the MinION reads due to the results on the simulated data. We then chose to compare the results to those obtained with Parallel-META 3. It had previously shown very high performance (Escobar-Zepeda et al. 2018) and was applicable to 16S rRNA data (see previous section). MetaG was run with its standard settings for MTX and Nanopore reads. Parallel-META 3 was run in its strictest alignment mode, which was three:

> PM-parallel-meta -t 4 -f F -e 3 -D B -r [input.fasta] -o [outpath]

The absolute taxon abundances given by both algorithms were transformed to relative abundances based on the individual amounts of matched reads. To the best of our knowledge, Cuscó and coworkers also focused on matched reads (Cuscó et al. 2018). The abundance of the unexpected taxa was the difference between the sum of the abundances of all expected taxa and a total of one. Plots were generated using R 3.4.4 and tidyverse 1.2.1.

## Data access

The source code of MetaG is available at https://github.com/IOB-Muenster/MetaG/tree/master/metag_src/ and the web service can be accessed at http://www.bioinformatics.uni-muenster.de/tools/metag/. Our simulated samples, customized databases and standard parameters can be accessed from https://github.com/IOB-Muenster/MetaG/tree/master/supplemental/files/query, https://github.com/IOB-Muenster/MetaG/tree/master/supplemental/files/db and https://github.com/IOB-Muenster/MetaG/tree/master/metag_src/install/files. A list of links to MetaG’s output for all presented samples is available on GitHub: https://github.com/IOB-Muenster/MetaG/blob/master/supplemental/files/query/README.md.

## Acknowledgements

The development of MetaG started as a master thesis, thus we are thankful for Dr. Francesco Catania’s supervision. We are grateful to Dr. Junya Yamagishi and participants of GRAID workshop for providing the MinION sequences from three pathogenic viral isolates. Dr. Victoria Shabardina helped to improve the concepts of MetaG in the early stages of algorithm development. We also thank Jonas Bohn and Reza Halabian, who were not afraid to test the early versions of the installation script and provided helpful comments for its improvement.

## Supplemental Materials

### Calculation of taxonomy confidence score

In the initial alignment stage of MetaG, each query read has zero, one or multiple database hits. For each read, MetaG assigns the taxonomy based on the highest number of hits. This is done at every taxonomic rank. However, the process is lacking statistics to evaluate “how good” a taxonomic assignment really is. In the following, the taxonomy confidence score will be explained at the example of a single query read. At a single rank, this hypothetical read has four potential taxonomies, called A, B, C and D (see Table S1). MetaG identifies taxon A as the taxon with the most hits for this read at this rank (see Table S1). The purpose of the following calculations is to show, whether A is the “true” taxon.

**Table S1:** Number of supporting hits and average e-value for four candidate taxa of a hypothetical query read. The resulting bin weights and the individual confidence are shown for each candidate.

| Taxon | Hit count | Average e-value | Bin weight | Individual confidence |
| --- | --- | --- | --- | --- |
| A | 4 | 0 | 4 | 16 |
| B | 3 | 0.2 | 2 | 6 |
| C | 1 | 0.4 | 1 | 1 |
| D | 1 | 0.6 | 0 | NA |

First, the candidate taxa are sorted by the alignment quality. This is first attempted by the arithmetic mean of the e-value (low to high), then by the arithmetic mean of the alignment score (high to low) or the alignment count for each taxon (high to low). In the resulting list, the first taxon has the best and the last one has the worst quality. The average e-values for the four taxa are given in Table S1. According to the quality (here the e-value), the taxa are grouped in a maximum of four bins. The first and the last taxon are automatically assigned to the first and the last bin, respectively. For the definitions of the other bins, we subtracted the worst from the best quality value and divided the result by three. Here, the result for the e-value is -0.2 (see Table S1). Bin two comprises of taxa which have a higher e-value than the best one and at max an e-value of best value - −0.2 = 0.2. The third bin ranges from the border of the second bin to quality values of at most best value - 2 * −0.2 = 0.4. All other taxa are assigned to the last bin. In our example, A is in the first bin, B is in the second bin, C and D are in the third and fourth bin, respectively (see Table S1).

Next, the bins are assigned with weights. Four for the first, two for the second, one for the third and zero for the last bin (see Table S1). For all consequent calculations the last bin is ignored. The individual count of hits for each taxon (see Table S1) is multiplied with its respective bin weight to receive the individual confidence. The individual confidences for the taxa are: A: 4 * 4 = 16, B: 3 * 2 = 6, C: 1 * 1 = 1 and D is ignored (see Table S1). The confidence of the taxon with the highest number of hits, A, is reduced by the individual confidences of all other taxa. This value is then divided by the total number of hits for all taxa, except for those in the last bin. Thus, the raw taxonomy confidence score for A is (16 – 6 – 1)/(4 + 3 + 1) which is ca. 1.13. For convenience, we adjust the range from −4 to 4 to the range between 0 and 1, bad and good confidence, respectively. Thus, the final taxonomy confidence score for A is ca. 0.64. There are two major exceptions from the calculations presented above: The taxon with the most hits will always get the highest taxonomy confidence score of one, if it is the only candidate taxon or if it has at least twice as many hits as the taxon with second most hits.

The confidence score for taxon A is multiplied by the confidence score for the taxon at the previous rank, if the currently considered rank is not the top one. This reflects the fact that a wrong assignment at higher ranks will bias the assignments at all lower ranks. Thus, assignments of taxa with lower confidences will lead to fast reduction of confidences at the lower ranks. If the current confidence value is lower than the cutoff, the confidence threshold that is set by the user will leave the current and all following taxa unassigned. Dependent on sequencing technology and database, we provide default taxonomy confidence scores in our standard parameters https://github.com/IOB-Muenster/MetaG/tree/master/metag_src/install/files.

### Building of databases

For the most part, databases were adapted to MetaG by using several custom scripts (https://github.com/IOB-Muenster/MetaG/tree/master/supplemental/scripts/db). Thus, later versions of the databases can be used without extensive manual labor. However, we made minor corrections to selected taxa. After modifying the database files, LAST databases were build using LASTdb (Frith et al. 2010):

> lastdb [dbPath][dbName] [db.fasta]

The databases used in our analyses can be found at https://github.com/IOB-Muenster/MetaG/tree/master/supplemental/files/db. The following sections will focus on building the databases from scratch.

### RDP

We downloaded the unaligned sequences for bacteria, archaea and fungi from the RDP website (https://rdp.cme.msu.edu/misc/resources.jsp). The fasta files were extracted and concatenated. Semicolons within the headers were replaces by commas. To get a taxonomy file for MetaG, we ran our makeRDPtax.pl script. For that, the db.fa must be located in the directory of the script. Apart from modifying the format, the script aimed to improve the taxonomic resolution by splitting the information for species and strain at the lowest taxonomic rank of RDP.

### MTX

We downloaded the Metaxa2.2 software from https://microbiology.se/software/metaxa2/. After extraction of the archive, we used blastdbcmd (Camacho et al. 2009) version 2.6.0+ to extract the BLAST databases for LSU and SSU sequences using:

> blastdbcmd -db [db]/blast -entry all -out [db].fa

Next, we merged the BLAST taxonomy files for the LSU and SSU sequences and customized these files. The steps included, but were not limited to, removing entries with missing lower ranks, splitting species and strain name and forcing the taxonomy to seven ranks. The latter was necessary, as there was no common number of ranks for all entries. Thus, we always took the first three and last four ranks. The complete set of changes can be made by running makeMTXtax.sh on the taxonomy file MTX.txt in the script directory. The output was tax.MTX.txt.

Using makeMTXfasta.pl, we removed the fasta entries for records which had been removed from the taxonomy file. For that, the fasta file MTX.fa and the modified taxonomy file tax.MTX.txt needed to be located in the directory of the script. The output was out.MTX.fa.

### ICTV

We retrieved the ICTV VMR database as an excel spreadsheet from https://talk.ictvonline.org/taxonomy/vmr/m/vmr-file-repository. We replaced empty fields with NA and removed line breaks in the fields. The field separator was set to tab and the sheet with the taxonomy data was saved in CSV format as VMR.csv. The script makeICTVvmr_db.pl must be located in the same directory. It queries the NCBI API to get the fasta sequences for all GenBank (Benson et al. 2018) IDs belonging to the database records. Viral taxa frequently had multiple GenBank IDs. All were reported, if they could be retrieved. The output was separated in three files holding the taxonomy, the fasta sequences and the pathogen information. These files were located in the directory of the script and were tax.VMR.txt, VMR.fa and patho.VMR.txt, respectively.

### PATRIC

In order to build a pathogen database for non-viral samples in MetaG, we obtained the genome_lineage and genome_metadata files from ftp://ftp.patricbrc.org/RELEASE_NOTES/. We modified the genome_metadata and removed all entries without a human host:

> awk -F ‘\t’ ‘{if ($46 ∼ /^Human|^Homo/) {print $1”\t”$46”\t”$53}}’ genome_metadata

The output was saved as patricHuman.txt. We then created our customized PATRIC database using the makePATRIC.pl script. This expected the genome_lineage and patricHuman.txt files to be located in the script directory. The output was patho.PATRIC.txt in the script directory.

## Training of MetaG

To find the best parameters for MetaG, we aligned our viral samples and the simulated marker gene samples to ICTV and to both MTX and RDP, respectively, using LAST. We kept these LAST alignments and created additional filtered alignments using LAST-SPLIT at -m 0.80, 0.85, 0.90, 0.95. Using these alignments, we simulated all possible combinations of the e-value, alignment score and confidence cutoff in the range from −40 to 10, 0 to 1 and 0 to 1, respectively, using custom scripts. These scripts reported the performance of each parameter combination and alignment type. For the viral samples, this was done based on the observed versus expected abundance of each taxon (trainKnownAbund.pl). For the simulated marker samples, the MCC for all identifications at genus level was used to avoid nomenclature clashes between the databases (trainKnownOrigin.pl). Detailed instructions how to run the scripts could be found in the help message of each script. Both scripts needed the CPAN (https://www.cpan.org/) modules Algorithm::Loops and Parallel::ForkManager. The scripts had to be placed in the same directory as metag.sh and the module directory of MetaG. Both scripts are available at https://github.com/IOB-Muenster/MetaG/tree/master/supplemental/scripts/train.

We manually examined the output and defined the standard parameters. We were looking for the strictest combination of e-value, alignment score and confidence cutoff which showed the highest performance. However, if the performance was equal over a wide range of parameters, we choose relaxed parameters from this subset. Performing LAST-SPLIT at -m 0.95 was preferred over using the native alignment, if the performance differences were negligible. This was done to improve the overall computations, as filtered alignments were often significantly smaller than the unfiltered alignments.

